# A series of terribly unfortunate events: How environment and infection synergized to cause the Kihansi spray toad extinction

**DOI:** 10.1101/2021.11.10.468118

**Authors:** Thomas R Sewell, Lucy van Dorp, Pria N Ghosh, Claudia Wierzbicki, Cristian Caroe, John V Lyakurwa, Elena Tonelli, Andrew Bowkett, Stuart Marsden, Andrew A Cunningham, Trenton W J Garner, Thomas P Gilbert, Ché Weldon, Matthew C Fisher

## Abstract

Outbreaks of emerging infectious diseases are trained by local biotic and abiotic factors, with host declines occurring when conditions favour the pathogen. Extinction of the Tanzanian Kihansi spray toad (*Nectophrynoides asperginis*) in 2004 was contemporaneous with the construction of a dam, implicating habitat modification in the loss of this species. However, high burdens of a globally emerging infection, *Batrachochytrium dendrobatidis* (*Bd*) were synchronously observed implicating infectious disease in this toads extinction. Here, by shotgun sequencing skin DNA from archived toad mortalities and assembling chytrid mitogenomes, we prove this outbreak was caused by the *Bd*CAPE lineage and not the panzootic lineage *Bd*GPL that is widely associated with global amphibian extinctions. Molecular dating showed an invasion of *Bd*CAPE across Southern Africa overlapping with the timing of the extinction event. However, post-outbreak surveillance of conspecific species inhabiting this mountainous region showed widespread infection by *Bd*CAPE yet no signs of amphibian ill-health or species decline. Our findings show that despite efforts to mitigate the environmental impact caused by dams construction, invasion of the pathogen ultimately led to the loss of the Kihansi spray toad; a synergism between emerging infectious disease and environmental change that likely heralds wider negative impacts on biodiversity in the Anthropocene.

## Main

Worldwide, amphibian populations are in steep decline being increasingly threatened by anthropogenic habitat alteration, climate change and the spread of infectious disease ^1–6^. Three decades of research on the disease chytridiomycosis and its causative pathogens, chytrid fungi in the genus *Batrachochytrium*, has proven the proximate link between these emerging infections and global amphibian declines ^2,3,7^. The epidemiology of these multi-host pathogens of ectothermic amphibians is extraordinarily complex with diverse biotic and abiotic factors known to modify the outcome of host-pathogen interaction post invasion ^2^. These factors include not only the suite of environmental variables that comprise the host amphibians fundamental ecological niche ^8,9^, but also the genotype of the invading pathogen itself ^10^. Accordingly, multiple lines of evidence are needed in order to establish causal associations between *Bd* infection and species declines or extinctions. However, owing to the intrinsic difficulty of working with wildlife alongside the understudied nature of many wildlife diseases, causal epidemiological inference is usually conducted *post hoc* and is qualitative rather than quantitative ^3,11,12^. This problem is compounded by the need to rely on the confirmation of amphibian chytridiomycosis along with *Bd* DNA in archived museum specimens ^13,14^. Yet, to date, the molecular detection of *Bd* from archived specimens has relied on either quantitative PCR or ITS-2 metabarcoding, neither of which provide reliable phylogenetic information on pathogen genotype or phylodynamic processes ^10,15^.

A striking example of an amphibian species extinction is that of the Kihansi spray toad (*Nectophrynoides asperginis*), an ovoviviparous bufonid, once micro-endemic to the Kihansi Gorge in the Udzungwa Mountains of Tanzania ^16–18^. This species had a spatially restricted population within the spray wetlands associated with waterfalls in the gorge ^16^. In 2000, the toads’ habitat experienced a ten-fold reduction in water flow due to the construction of the Lower Kihansi Hydropower Project, following which the spray toad population sharply declined to less than 2,000 individuals by March 2001 ^16,19^. Conservation measures to restore the wetland’s ecologically important habitat initially appeared successful, with moderate habitat regeneration and rapid recovery to almost 18,000 toads by June 2003 ^16,20^. However, the spray toad population entered a catastrophic and enigmatic decline soon after the 2003 survey and in 2009 the species was officially declared extinct in the wild ^17^. Factors connected to the Lower Kihansi Hydroelectric Project, such as a modification of wetland size and composition, ineffective mitigation measures and the accidental release of contaminated sediments were considered principal drivers of habitat alteration and the consequent decline of the *N. asperginis* population ^16,21,22^.

Evidence of *Batrachochytrium dendrobatidis* (*Bd*) infection in Kihansi spray toads was first recorded during the terminal decline of the wild population ^20^. *Post hoc* examination of archived spray toads, collected from the Kihansi Gorge between 1996 and 2003, established that *Bd* was not present until the start of the ultimate population decline, at which time histopathological examination detected severe burdens of infection that were consistent with those known to be lethal ^22^. To more fully understand the role that *Bd* played in the extinction event, we accessed the last two ethanol preserved Kihansi spray toads that had been collected at the time of the 2003 population crash. Using this archived fixed tissue, we applied sequencing protocols developed for the analysis of ancient DNA to shotgun sequence amphibian skin biopsies to determine whether we could assemble the pathogens genome ^23^. By combining our DNA reads with those from pre-existing *Bd* genomic datasets we were able to successfully assemble complete 178 kbp *Bd* mitochondrial genomes and in doing so demonstrate the feasibility of using archived specimens for phylodynamic analyses. Such an approach holds promise to provide greater insight into the factors underlying population declines and extinctions of other amphibians globally.

### Analysis and Results

Our genetic analyses found that the two Kihansi spray toads collected during the terminal population decline were infected with *Bd*CAPE, the endemic African lineage of the *Bd* ^10^. This finding, coupled with previous histological and molecular diagnoses of chytridiomycosis ^22^, provide the first evidence of a *Bd*CAPE associated extinction event in the wild.

Initial exploratory analysis using a newly-developed lineage specific diagnostic assay ^15^ found that the two Kihansi spray toad specimens showed triplicated qPCR amplification for the *Bd*CAPE lineage and no amplification for *Bd*GPL. Subsequently, we were able to generate enough *Bd* DNA reads from archived amphibian tissue samples to assemble the *Bd* mitochondrial genome. During the analysis, we controlled for low read depth by using a joint variant calling approach (GATK Best Practices) which utilised previously sequenced *Bd* isolates representing the known global diversity of this pathogen (Table S1) ^10^. This approach allowed us to leverage information from lineage-specific variants across a cohort of global isolates sequenced at a much greater depth, and in doing so providing us with increased confidence in variants that were called at a low depth in the two Kihansi samples. Samples AC040803_A and AC290703_A2 generated 252,950,676 and 294,104,776 Illumina sequencing reads, respectively; of which, 28,745 (AC040803_A) and 15,343 (AC290703_A2) reads mapped successfully to the *Bd* mitochondrial reference assembly of isolate JEL423 (Table S2). Average depth of coverage when aligned to the reference mitochondrial assembly was 8.97x and 4.65x, respectively (Fig S1). At a depth of >2’, the AC040803_A alignment had a total of 160,904 bases covered (92.41%) and the AC290703_A2 alignment had 142,867 bases covered (82.05%). The average allele balance across both samples was 0.98, suggesting a single genotype associated with each infected toad.

Phylogenetic analysis of 1,205 mitochondrial variants revealed that the two Kihansi samples were genetically highly similar to one another, differing by only 46 SNPs. These genotypes were also closely related to previously sequenced isolates of *Bd*CAPE, clustering closely with two chytridiomycosis-associated isolates, DB8-4 (Drakensburg, South Africa) and TF5a1 (Torrent des Ferrerets, Mallorca). All five deeply diverged *Bd* lineages are represented in the phylogeny, with globally high bootstrap support and therefore confidence in the placement of AC040803_A and AC290703_A2 in the *Bd*CAPE lineage (Fig 1A). In addition, a maximum-likelihood genetic clustering approach based on 724 variants, together with a principal component analysis, further supported the genetic relatedness of the two Kihansi samples with that of contemporary *Bd*CAPE isolates recently collected in Southern Africa (Fig 1B). Wards clustering designated AC040803_A and AC290703_A2 as a member of the same cluster as other *Bd*CAPE isolates and a PCA partitions diversity of other *Bd*CAPE isolates away from the *Bd*GPL cluster (Fig 1C).

**Fig. 1.**
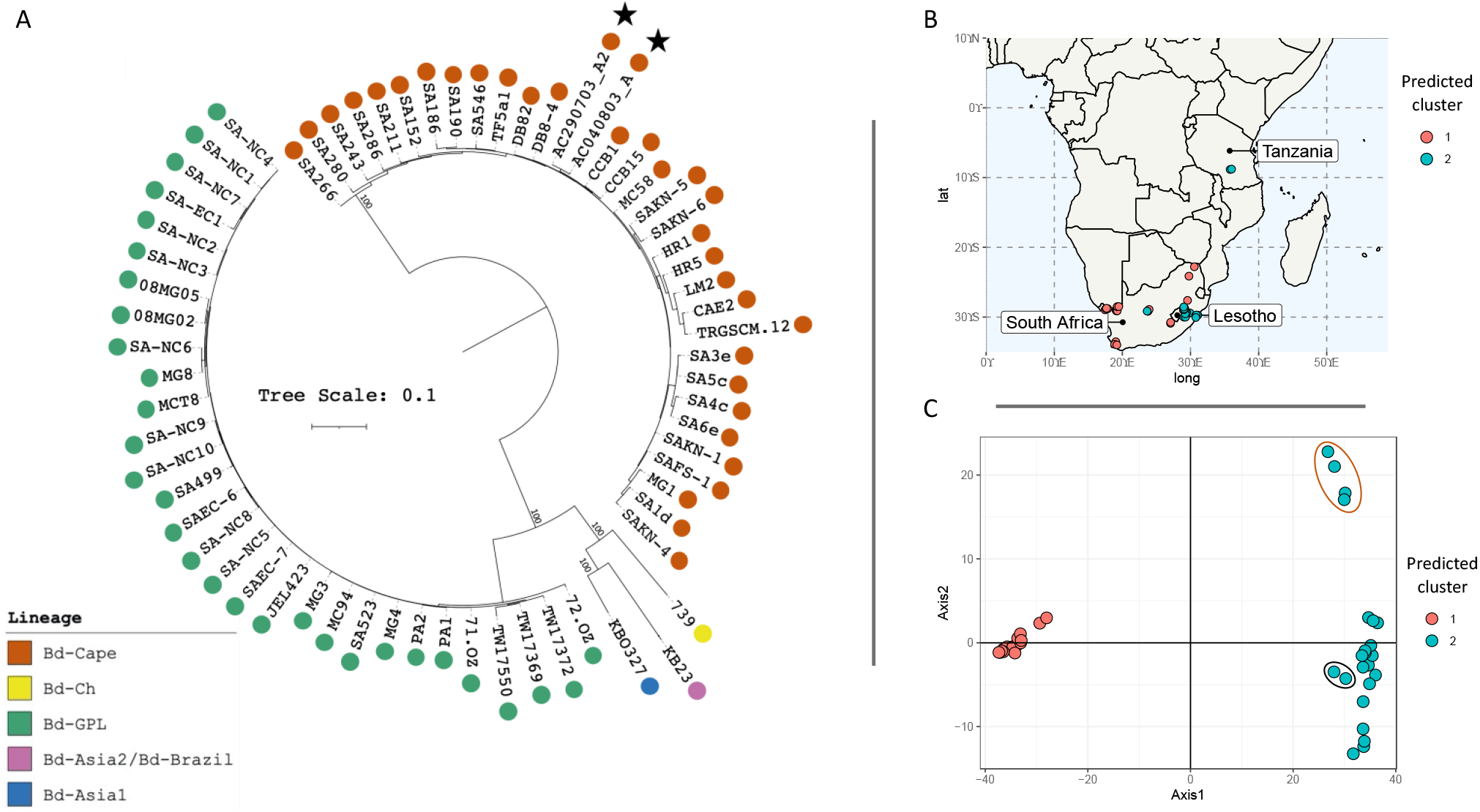
**A)** A midpoint rooted maximum likelihood (ML) tree constructed from 1,205 high-quality SNPs within the 178-kbp mitochondrial genome. The phylogeny was generated using RAxML utilizing the generalized time reversible (GTR) model and category (CAT) rate approximation. The two Kihansi archival samples are highlighted with stars. The five major lineages of *Bd* are denoted by tip color with good bootstrap support between divergence events. **B)** Map of southern Africa indicating the location of *Bd*CAPE and *Bd*GPL isolates considered for maximum-likelihood genetic clustering. **C)** Principal component analysis (PC 1-2) based on mitochondrial SNPs (45 individuals, 724 SNPs), generated from WGS of cultured South African *Bd* isolates plus the two Kihansi archival samples (circled black). Colours are based on predicted clustering generated using snapclust in R, which fully resolved both *Bd*GPL and *Bd*CAPE lineages. Four Pinetown isolates that harbored regions of elevated SNP density are circled in orange.

Our current understanding of the epidemiology and virulence of *Bd*CAPE is limited ^24,25^. Experimental studies have shown that *Bd*CAPE can cause lethal chytridiomycosis and is more frequently associated with chytridiomycosis in wild species than is seen for *Bd*ASIA-1, *Bd*ASIA-2/BRAZIL or *Bd*ASIA3 ^2,10^. Moreover, *Bd*CAPE is known to have caused a population decline following its introduction into the Balearic Island population of Mallorcan midwife toads (*Alytes muletensis*) ^8,26,27^. Despite this, there remains a general perception that widely globalized *Bd*GPL is the only lineage of conservation concern. Although *Bd*GPL has been documented in many more declines than *Bd*CAPE, and has a much broader distribution ^3,10^, recent findings demonstrate that the *Bd*CAPE lineage is more widely distributed than previously thought ^28^. Records of *Bd*CAPE from Europe and Central America most likely represent introductions from southern or western Africa, where the majority of *Bd*CAPE observations have occurred and where genetic diversity appears to be high ^10,28^. Our evidence of *Bd*CAPE causing severe chytridiomycosis in Kihansi spray toads during their decline expands the range of this lineage into a new area and highlights the threat posed to amphibian hosts by other, non-*Bd*GPL lineages.

The two genotypes infecting the Kihansi spray toads were closely related to previously sequenced *Bd*CAPE isolates, both known to cause amphibian chytridiomycosis or symptoms associated with chytridiomycosis. Isolate TF5a1 was collected from the Torrent de Ferrerets gorge during the Mallorcan midwife toad outbreak of chytridiomycosis, which was first witnessed in 2008 ^24,27^. This isolate has been shown experimentally to cause chytridiomycosis in the common toad (*Bufo bufo*) in which it was hypovirulent when compared to a *Bd*GPL ^24^. Isolate DB8-4 was collected atop the Drakensburg portion of the Great Escarpment bordering South Africa and Lesotho, from a population of Phofung river frogs (*Amietia hymenopus*) that has been anecdotally observed to show clinical signs of chytridiomycosis ^29^. In light of these observations, and the histological evidence published by Weldon et al. ^22^, it would appear that the genotype found infecting the Kihansi spray toads was typical of *Bd*CAPE genotypes circulating within Southern Africa, and that are known to be virulent in nature.

To assess the timing of emergence of *Bd*CAPE we first confirmed the presence of significant temporal signal in the alignment (Fig S2) once confounding SNPs in four Pinetown isolates were excluded (Fig S3). We identified a positive correlation between root-to-tip phylogenetic distance and time of sampling which was significant following date randomization (Fig S2A-C). Subsequently, bayesian tip-dating was applied in BEAST2 ^30^, testing various plausible demographic and clock models run until model convergence occurred (Table S3A). Following nested path sampling, the model with the highest marginal likelihood was found to be a relaxed clock with a coalescent constant population prior (Table S3B). The mean mitochondrial substitution rate was estimated as 2 × 10^−5^ substitutions per site per year [95% highest posterior density (HPD), 6 × 10^−6^ to 3.5 × 10^−5^] (Fig S4A). This rate was marginally faster than that previously estimated for the *Bd*GPL mitolineages, at 1.01 × 10^−6^ substitutions per site per year ^10^. Our analysis supports a recent tMRCA of sampled diversity to between 14 and 67 years ago (mean of 26 years since the most recent sampling date), pointing to an African introduction of *BdCAPE* between 1944 and 2003, with tMRCA of 1991 (Fig 2A-B). Regardless of model specification, all estimates of posterior density encompass this time period (Fig S5A-D). Importantly, our estimated age of the common ancestor of African *Bd*CAPE suggests that the lineage invaded the Kihansi gorge region around the time that the hydropower dam was constructed. In this case, would the Kihansi Spray toads have succumbed to chytridiomycosis without the presence of the substantial environmental perturbation caused by the damming of the gorge?

**Fig. 2.**
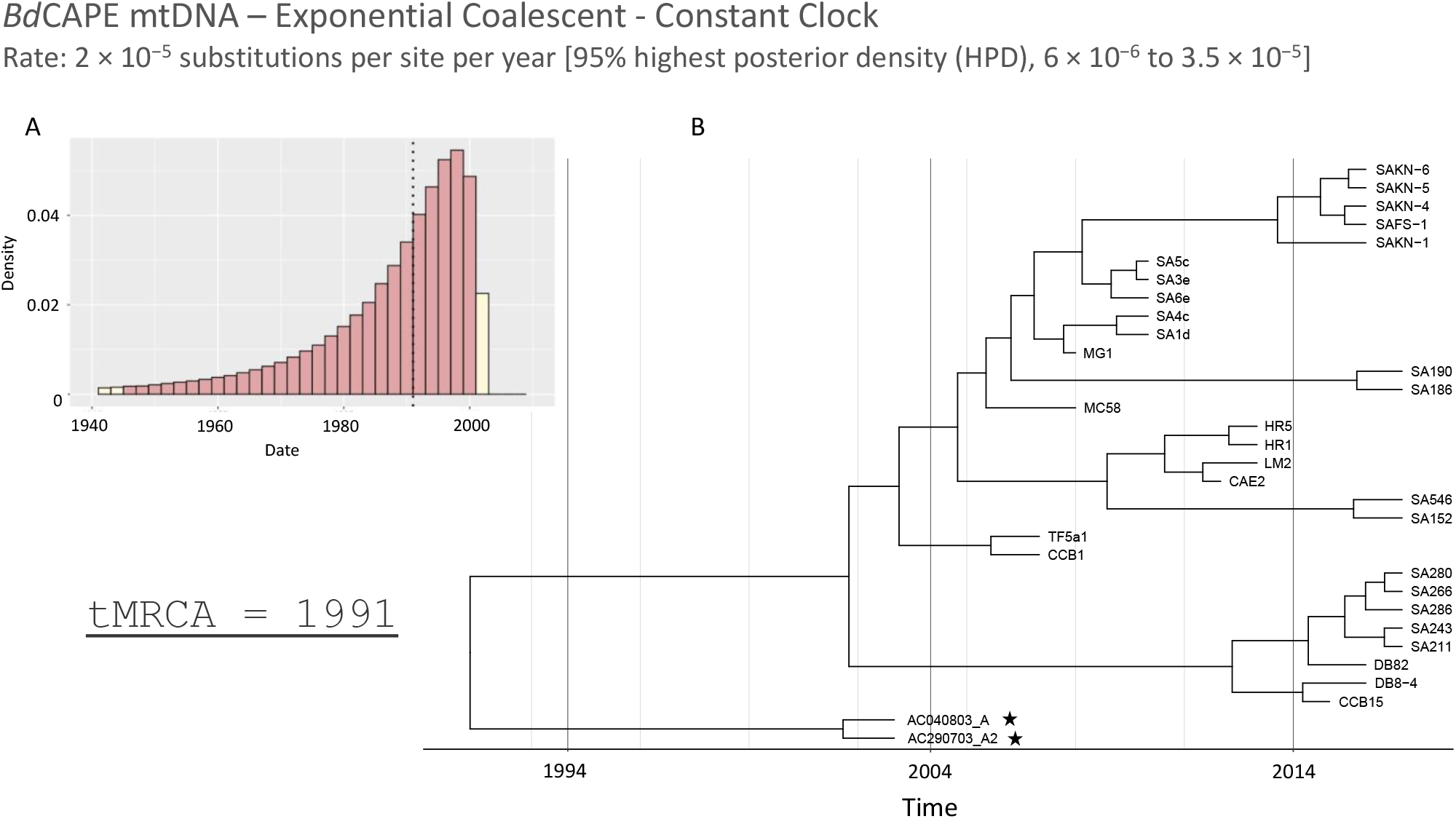
Bayesian tip-dating analysis of *Bd*CAPE mitochondrial diversity in Africa and with node calibrations estimated using BEAST2. The mean mitochondrial substitution rate was 2 × 10^−5^ substitutions per site per year [95% HPD, 6 × 10^−6^ to 3.5 × 10^−5^] **A)** Posterior distribution for tMRCA (tree height). Red bars indicate 95% HPD intervals of 14 and 67 years since most recent sampling date of 2017. Dotted black line indicates the mean tMRCA of sampled diversity to 1991 (26 years since most recent sampling date of 2017). B) Maximum clade credibility phylogeny estimated using median node heights. Yellow bars indicate the 95% HPD for node heights. The two Kihansi archival samples are highlighted with a star.

To answer this question, we undertook wider surveys of *Bd* across the landscape that was home to the spray toads, the Udzungwa mountain range. We took non-invasive skin swab samples from 24 amphibian species across a distance of ~30 km (Fig 3). Of the 60 swab samples, 15 (25%) returned positives for *Bd*, with a GE count of between 0.01 and 71.1. These results echo that of a previous and more extensive *Bd* survey in the region that was completed in 2006, just after the spray toad’s terminal population decline ^31^. Of the five *Bd* positive samples that yielded a lineage-specific result, all were confirmed as *Bd*CAPE (Table S4). All amphibian species that were positive for *Bd* showed no signs of disease and or population declines during the three-year survey period. Importantly, *Nectophrynoides tornieri* or Tornier’s viviparous toad (Fig. 3), a close relative of the Kihansi spray toad, was found to be infected by *Bd*CAPE infection yet displayed no evidence of chytridiomycosis or population decline.

**Fig. 3.**
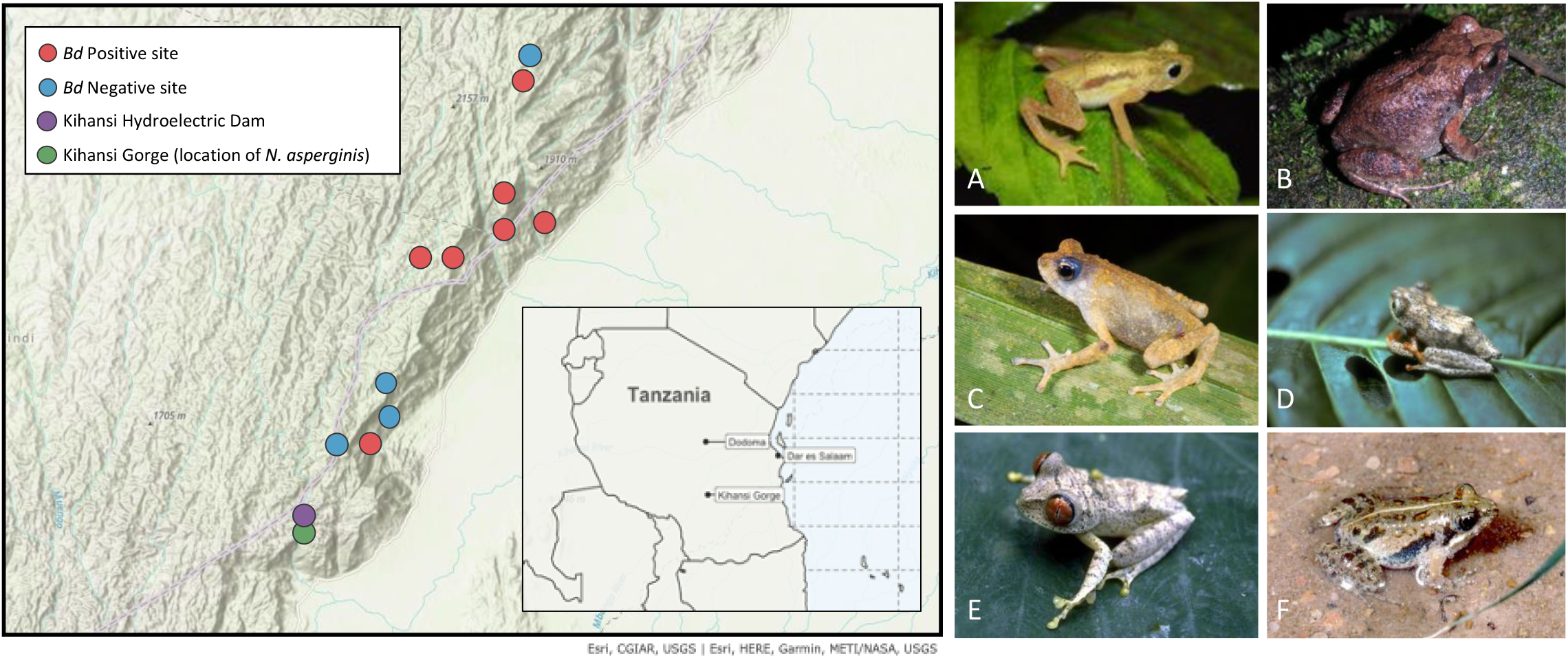
A map of the Udzungwa Mountain region in Tanzania outlining the locations of the qPCR swab survey. Positive swab sites are marked in red and negative sites marked as blue. The location of the Kihansi power station is marked in purple and the Kihansi spray toad site in green. Small insert map of Tanzania shows the location of the Kihansi gorge relative to major cities. Photographs of the Kihansi spray toad and amphibian species that tested positive for *Bd*CAPE in 2003 and during the qPCR swab survey a) *Nectophrynoides asperginis* b) *Arthroleptis stenodactylus* c) *Nectophrynoides tornieri* d) *Hyperolius kihangensis* e) *Leptopelis parkeri* f) *Phrynobatrachus sp* – Photography credits: Robert C. Drewes; Martin Pickersgill; Stephen Zozaya; John V Lyakurwa; Elena Tonelli

## Discussion

Through deep sequencing archived *Bd* infected toads as they entered their terminal decline in the wild, we were able to form a nuanced understanding of the epidemiology of this extinction-by-infection event. The Kihansi spray toad population was only discovered in 1996 and began to decline soon thereafter ^16,19,22^. Although retrospective designation of disease as a primary driver of population declines or extinctions is highly challenging ^11,12^, our analysis clearly identifies an outbreak of *Bd*CAPE as a proximate factor in the terminal decline of this species. Moreover, that other amphibian species including the conspecific *Nectophrynoides tornieri* in the Udzungwas are infected with *Bd*CAPE yet show no evidence of disease, is highly suggestive that ecological change wrought by damming the Kihansi gorge amplified the impact of the invading pathogen; parallel climate-disease interactions have been noted in other *Bd-* amphibian systems ^9,32^. Importantly, that *Bd*CAPE is now known to occur widely, can cause lethal amphibian chytridiomycosis ^22,24,26,27^, and has now been associated with an amphibian extinction event, we clearly need to upgrade our assessment of the risk that this lineage poses to amphibian species more widely.

Using archived specimens to better understand the factors leading amphibian declines is an approach that holds promise in determining the drivers underlying species extinctions, and to better inform on historic zoonotic spillover events. Our results emphasize the threat that emerging infectious diseases have on small, isolated populations when in combination with factors that alter ecologically important habitats. In the scenario we present, this emerging pathogen appears to have synergised with other threats, with the population eventually becoming so small that extinction in the wild occurred where no one threat alone would otherwise have a lead to a terminal decline. That other amphibian species in the region endured no such impact supports our thesis that the invading pathogen was able to cause lethal infections on a host that terminally succumbed to a series of terribly unfortunate events.

## Methods

### Kihansi Spray Toad Sample Collection and DNA extraction

Two preserved specimens of Kihansi spray toad (*Nectophrynoides asperginis*) were provided by Prof Che Weldon of North-West University. The spray toads were originally collected dead from the Kihansi Gorge in the Udzungwa Mountains, Tanzania (−8.575, 35.851389) (Fig S6) between June and August 2003; this was during the spray toads terminal population decline ^22^. The samples were kept in ethanol until use and were processed in a UV-treated clean room (free from amplified PCR product and cultured DNA extracts). One foot from each Kihansi spray toad was removed and air dried for two hours. DNA was extracted using Qiagen DNEasy Blood and Tissue Kit (Section 2.6) according to the manufacturer instructions, including the recommended double elution step to maximize DNA yield.

### Swab sampling across the Udzungwa Mountains

Between November 2013 and March 2015, 60 adult and juvenile amphibians were swabbed for *Bd* in the Uzungwa Scarp Nature Forest Reserve, north-east of the Kihansi Gorge. Fine-tipped MW100 swabs (MWE, UK) were used to gently run the swab over the back, ventre and legs, ensuring the back legs and toes were well swabbed. All swabs were kept at environmental temperature (14-30°C) for the entire duration of field work. DNA was extracted directly from swabs in accordance with the *RACE* protocol and were processed in a dedicated pre-PCR clean room in a decontaminated and UV sterilised cabinet. (free from amplified PCR product and cultured DNA extracts). Swab tips were snapped off and placed directly into Safe-Lock Eppendorf tubes containing 0.03 to 0.04g of 0.5mm silica homogenisation beads and 60μl of Prepman Ultra (ThermoFisher Scientific, Massachusetts, USA). Samples were bead-beated for 45 seconds at 30Hz in a Qiagen TissueLyser II (Qiagen, Venlo, Netherlands) before being centrifuged at 14,500rpm for 30 seconds, a process that was repeated twice in total. Samples were then incubated at 100.°C for 10 minutes, cooled for two minutes, then centrifuged again at 14,500rpm for three minutes. The supernatant was collected and transferred to a fresh 0.5ml or 1.5ml Safe-Lock Eppendorf tube.

### qPCR assay on Kihansi samples and Udzungwa Mountains survey swabs

Due to a need to conserve DNA available for shotgun sequencing, Kihansi spray toad DNA extracts were not tested with the pan-lineage *Bd* diagnostic ^33^ but moved straight onto a lineage-specific qPCR assay ^15^, that distinguishes between *Bd*CAPE and *Bd*GPL lineages. Two of the four archived samples that showed the highest number of *Bd* genomic equivalent score (GE), AC040803_A and AC290703_A2, were then shotgun-sequenced to confirm the lineage diagnosis and generate genotypic data for downstream analyses. Swab samples collected in the Udzungwa Mountains, Tanzania were processed according to the pan-lineage *Bd* qPCR diagnostic ^33^. Swabs that had a >1 GE, were then processed using the lineage-specific qPCR assay ^15^.

### Library preparation and sequencing of Kihansi Spray Toad samples

Libraries were built using the Nebnext Ultra II FS library kit (New England Biolabs, Ipswich, ma, US) with an average target fragment length of 300-700bp. Adaptor-ligated libraries were purified with 1x volume of SPRI beads ^34^. qPCR was carried out to estimate amplification cycle numbers for the indexing PCR. Libraries were subsequently amplified using NEBs Q5 2x mastermix for 10 cycles. Amplified libraries were purified with SPRIbeads as before and purified libraries were analysed on an Agilent 2100 Bioanalyzer. Pooled samples were sequenced on two lanes on an Illumina HiSeq 4000 instrument in single read mode for 80 cycles.

### Read alignment and variant identification

Genome Analysis Toolkit (GATK) v3.6 was used to call variant and reference nucleotides from two subsets of *Bd* mitochondrial alignments; firstly, a collection of previously sequenced *Bd* isolates representing five deeply diverged *Bd*-lineages for phylogenetic analysis ^10^; secondly, a novel collection of WGS *Bd*GPL and *Bd*CAPE isolates collected in South Africa for *de novo* genetic clustering. The two Kihansi samples, AC040803_A and AC290703_A2, were included in each set of alignments for comparison. All sequenced reads were trimmed using cutadapt v1.9.1 to remove adapter sequences and low-quality ends (Phred score <20). BWA v0.7.8 was used to align the trimmed reads to the both the nuclear and mitochondrial reference assembly of the *Bd* isolate Jel423 (Broad), and samtools v1.3.1 was used to generate mapping statistics. Mitochondrial alignments were selected for subsequent analysis due to their percentage breadth of coverage. Prior to variant calling, alignments were pre-processed using AddOrReplaceReadGroup, to assign all reads per file to a single new read-group tag, and MarkDuplicates to identify duplicate reads. Variants were detected using HaplotypeCaller (GATK) in joint calling mode and with ploidy set to one. SelectVariants (GATK) was used to generate a SNP-only VCF file, and VariantFiltration was used to filter for high-quality variants (DP < 4 || MQ < 40.0 || QD < 2.0 || QUAL < 50).

### Phylogenetic and population genetic analysis

SNPs generated from the subset of samples used for phylogenetic analysis were concatenated into a multi-sample FASTA and converted into PHYLIP format. As the variants were identified simultaneously across all samples using joint calling mode in GATK (where genotype calls are generated at every site where any sample in the call set has evidence for variation), a representative curated alignment of 1,205 mitochondrial nucleotides was used for phylogenetic inference. A phylogenetic tree was generated using RAxML v8.2.9, employing the generalised time reversible (GTR) model and category (CAT) rate approximation. Phylogenetic uncertainty was assessed using 500 bootstrap iterations implemented via RAxML’s rapid bootstrapping mode. The consensus phylogeny was imported into R (*29*) and visualised with associated bootstrap support using ggtree (v1.16.6) ^36^.

A multi-sample VCF file (725 SNP variants), generated from the subset of samples used for maximum-likelihood genetic clustering, was imported into R using vcfR ^37^ and analysed using the snapclust function in the adegenet package ^38^. As prior lineage association of samples was known for all isolates excluding the two Kihansi samples, snapclust was supervised to discriminate two clusters (k=2; *Bd*-GPL and *Bd*-Cape). The Ward clustering algorithm was applied for group membership. Principal coordinate analysis (PCA) of mitochondrial variants was used to evaluate the clustering of isolates, which were coloured according to the group membership output generated by snapclust.

### Bayesian dating of *Bd*CAPE diversity in Africa and tMRCA estimations

Bayesian phylogenetic tip-dating was conducted in BEAST2 (v2.6.3) ^30^, using an alignment of 30 *Bd*CAPE isolates with recorded sampling years between 2008 and 2017, plus the two Kihansi archival samples that were collected in 2003 (a final alignment of 32 individuals and 674 mitochondrial variants). Clustering analysis of African BdCAPE isolates revealed a two-cluster split (Fig 1B), with four isolates from Pinetown, South Africa forming their own separate cluster away from the remaining *Bd*CAPE genotypes. To assess the SNPs underlying this clustering we ran Gubbins (v2.3.4) and Phandango (v1.3.0) ^39,40^ together with manual inspection of the alignment. Three putatively recombinant blocks were identified (A-C), distributed across all four Pinetown isolates and confirmed following visual check of the alignment (Fig S3). Removal of these regions of high SNP density resulted in a final alignment of 195 mitochondrial variants. A maximum-likelihood phylogeny was generated using iqtree (v1.6.12) with 100 bootstrap values to observe the effect pruning had on the resulting maximum likelihood tree topology ^41^. A reduction in the length of the terminal branches was observed in the Pinetown cluster and the placement of isolate SA4c within the diversity observed among other *Bd*CAPE isolates. The roottotip() function in BactDating was applied to identify the presence of a significant root-to-tip regression which was assessed for significance following 1000 random permutations of sampling dates ^42^. Phylostems was also applied to visualize the extent of temporal correlation at each node in the phylogeny ^43^. Both tools supported a measurably evolving alignment, justifying application of phylogenetic tip-calibration using all the southern African *Bd*CAPE isolates in our collection.

Using BEAST2 we initially applied a model averaging approach via the package bModelTest (v1.2.1) to identify the best supported site model for the dataset ^30,44^. The two models with highest posterior support were 121324 - 25% posterior support (TN93) and 123423 - 22.97% posterior support (TIM). Given there was no strong preference for these or any other substitution models we proceeded with a model averaging approach. We then applied both relaxed and strict clock models across one of four possible demographic priors: Coalescent Constant Population, Coalescent Exponential Population, Coalescent Bayesian Skyline and Coalescent Extended Bayesian Skyline. BEAUti (v2.6.3) was used throughout to generate the xml input files ^30^. To ensure an appropriate rate calculation, the xml files were manually edited to include all the counts of invariant A, T, C & G sites. The MCMC chain length for all runs was set to 5×10^8^ and the trace log was set to record every 1000 samples, with convergence confirmed by an Effective Sampling Space (ESS) >200 for all parameters and visual inspection of the MCMC traces. In addition, all models were run without the data – sampling from the prior – to confirm the posteriors estimated were not being driven by the choice of priors. To select the most appropriate clock and demographic model combination for our data, we applied the nested sampling function implemented in the NS package (v1.0.3) ^45^. Bayes Factors (BF) were generated through pairwise comparison of the estimate marginal likelihoods and interpretation of the BF ranges was made with accordance to Kass and Raftery, 1995 ^46^ (Table S4B). All log output files were processed in R using the beastio and bdskytools packages (https://github.com/laduplessis) with the first 10 % of the chain discarded as burn-in. Posterior distributions for each model were plotted using ggplot2 ^47^. To summarise the tree topology, a maximum clade credibility tree was generated using Tree Annotator (v2.6.3) specifying a 10% burn in percentage and considering the median value of node heights. The time calibrated phylogeny was visualised and annotated using ggtree in R ^36^.

## Acknowledgments

DNA sequencing was carried out at the Danish National High-throughout DNA Sequencing Centre in Copenhagen, Denmark. Samples from the Udzungwa Mountains were collected with permission from the Tanzanian Commission for Science and Technology, Tanzania Wildlife Research Institute and Tanzania Forest Services Agency (Permit numbers: TFS./106/552/01/08 and TFS/KL/167(1)291/15). ET research permits for the two field seasons Nov 2013-March 2014 and Nov. 2014 – March 2015: COSTECH permits No. 2013-333-ER-2008-157 and 2014-338-ER-2008-157. The two Kihansi spray toad samples were collected under the auspices of LKEMP (Lower Kihansi Environmental Management Project).

## Author contributions

TRS, PNG, TWJG, TPG, AAC, CW and MCF conceived and designed the study. PNG, CWi, CC, JVL, ET and CWe collected the data. TRS, LvD and PG analyzed the data. TRS and PNG wrote the manuscript. TRS, LvD, PNG, CWi, CC, ET, AEB, SM, AAC, TPG, TWJG, CWi and MCF discussed the results and commented on the manuscript.

## Data and materials availability

Shotgun sequencing data have been deposited in the National Center for Biotechnology Information (NCBI) Sequence Read Archive (SRA). All sequences are available from NCBI BioSample accession: SAMN20599532. All data analysis scripts have been made available at github.com/swelbo/kihansi_paper.

