## Supplementary documents for "A series of terribly unfortunate events: How environment and infection synergized to cause the Kihansi spray toad extinction"

### Supplementary Materials:

Figures S1-S6

Tables S1-S4

Table S1: Table of *Batrachochytrium dendrobatidis* (*Bd*) isolates used during this study, plus the two Kihansi samples. Applies to the analysis of population structure to determine Kihansi sample lineage grouping (Fig 1) and the phylogenetic dating using BEAST2 (Fig 2).

| Isolate name | Country of isolation | latitude | longitude | Host species | Host Family | Lineage | Year of isolation | Used in population structure? | Used in phylogentic dating? |
|---|---|---|---|---|---|---|---|---|---|
| SA-NC4 | South Africa | -28.91356 | 19.0014167 | *NA* | Pyxicephalidae | GPL | 2015 | Yes | |
| SA-NC1 | South Africa | -28.91356 | 19.0014167 | *NA* | Pyxicephalidae | GPL | 2015 | Yes | |
| SA-NC7 | South Africa | -28.73725 | 19.299222 | *NA* | Pyxicephalidae | GPL | 2015 | Yes | |
| SA-EC1 | South Africa | -30.598095 | 29.894511 | *NA* | Pyxicephalidae | GPL | 2015 | Yes | |
| SA-NC2 | South Africa | -28.91356 | 19.0014167 | *NA* | Pyxicephalidae | GPL | 2015 | Yes | |
| SA-NC3 | South Africa | -28.91356 | 19.0014167 | *NA* | Pyxicephalidae | GPL | 2015 | Yes | |
| 08MG02 | South Africa | -28.76268 | 28.91794 | *Amietia vertebralis* | Pyxicephalidae | GPL | 2008 | Yes | |
| 08MG05 | South Africa | -34.093316 | 18.42167 | *Amietia fuscigula* | Pyxicephalidae | GPL | 2008 | Yes | |
| SA-NC6 | South Africa | -28.73725 | 19.299222 | *NA* | Pyxicephalidae | GPL | 2015 | Yes | |
| MG8 | South Africa | -23.85 | 30.033333 | *Amietia angolensis* | Pyxicephalidae | GPL | 2008 | Yes | |
| MCT8 | South Africa | -34.093316 | 18.42167 | *Amietia fuscigula* | Pyxicephalidae | GPL | 2008 | Yes | |
| SA-NC9 | South Africa | -28.69598 | 17.597361 | *NA* | Pyxicephalidae | GPL | 2015 | Yes | |
| SA-NC10 | South Africa | -28.69598 | 17.597361 | *NA* | Pyxicephalidae | GPL | 2015 | Yes | |
| SA499 | South Africa | -29.08682 | 23.79704 | *Amietia* | Pyxicephalidae | GPL | 2017 | Yes | |
| SAEC-6 | South Africa | -30.72189 | 26.9064555 | *NA* | Pyxicephalidae | GPL | 2016 | Yes | |
| SA523 | South Africa | -29.08682 | 23.79704 | *Amietia* | Pyxicephalidae | GPL | 2017 | Yes | |
| SA-NC8 | South Africa | -28.69598 | 17.597361 | *NA* | Pyxicephalidae | GPL | 2015 | Yes | |
| SA-NC5 | South Africa | -28.91356 | 19.0014167 | *NA* | Pyxicephalidae | GPL | 2015 | Yes | |
| SAEC-7 | South Africa | -30.72189 | 26.9064555 | *NA* | Pyxicephalidae | GPL | 2016 | Yes | |
| JEL423 | Panama | 8.5833 | -82.5333 | *Agalychnis lemur* | Phyllomedusidae | GPL | 2004 | Yes | |
| MG3 | South Africa | -27.682836 | 29.57863 | *Amietia angolensis* | Pyxicephalidae | GPL | 2008 | Yes | |
| MC94 | South Africa | -22.901675 | 30.696089 | *Xenopus laevis* | Pipidae | GPL | 2008 | Yes | |
| MG4 | South Africa | -34.093316 | 18.42167 | *Amietia fuscigula* | Pyxicephalidae | GPL | 2008 | Yes | |
| PA1 | Chile | -33.44713044 | -70.65281852 | *Calyptocephalella gayi* | Calyptocephalellidae | GPL | 2015 | Yes | |
| PA2 | Chile | -33.44713044 | -70.65281852 | *Calyptocephalella gayi* | Calyptocephalellidae | GPL | 2015 | Yes | |
| 71.OZ | Australia | -31.90714775 | 115.7937948 | *Litoria Moorei* | Pelodryadidae | GPL | 2016 | Yes | |
| TW17550 | Taiwan | 23.9497 | 120.8539 | *Buergeria japonica* | Rhacophoridae | GPL | 2017 | Yes | |
| TW17369 | Taiwan | 23.8897 | 120.7565 | *Hylarana latouchii* | Ranidae | GPL | 2017 | Yes | |
| TW17372 | Taiwan | 23.8897 | 120.7565 | *Hylarana latouchii* | Ranidae | GPL | 2017 | Yes | |
| 72.OZ | Australia | -31.54236573 | 115.6823222 | *Litoria Moorei* | Pelodryadidae | GPL | 2016 | Yes | |
| 739 | Switzerland | 47.30691163 | 8.498080357 | *Alytes obstetricans* | Alytidae | CH | 2007 | Yes | |
| KBO_327 | South Korea | 38.1298 | 127.7657 | *Bombina orientalis* | Bombinatoridae | ASIA-1 | 2014 | Yes | |
| KB23 | South Korea | 38.1298 | 127.7657 | *Rana catesbeianus* | Ranidae | ASIA-2 | 2014 | Yes | |
| SAKN-4 | South Africa | -28.71981 | 28.923611 | *NA* | Heleophrynidae | CAPE | 2016 | Yes | Yes |
| SA1d | South Africa | -28.71986 | 28.92375 | *Hadromophryne natalensis* | Heleophrynidae | CAPE | 2010 | Yes | Yes |
| MG1 | South Africa | -28.844605 | 29.054547 | *Amietia vertebralis* | Pyxicephalidae | CAPE | 2008 | Yes | Yes |
| SAFS-1 | South Africa | -28.75064 | 28.871361 | *NA* | Pyxicephalidae | CAPE | 2016 | Yes | Yes |
| SAKN-1 | South Africa | -28.75231 | 28.894361 | *NA* | Pyxicephalidae | CAPE | 2016 | Yes | Yes |
| SA3e | South Africa | -29.02242 | 30.58106 | *Amietia angolensis* | Pyxicephalidae | CAPE | 2010 | Yes | Yes |
| SA4c | South Africa | -29.02242 | 30.58106 | *Amietia angolensis* | Pyxicephalidae | CAPE | 2010 | Yes | Yes |
| SA5c | South Africa | -29.02242 | 30.58106 | *Amietia angolensis* | Pyxicephalidae | CAPE | 2010 | Yes | Yes |
| SA6e | South Africa | -29.02242 | 30.58106 | *NA* | Pyxicephalidae | CAPE | 2010 | Yes | Yes |
| LM2 | UK | 51.53659 | -0.153415 | *Leptopelis rufus* | Arthroleptidae | CAPE | 2013 | Yes | |
| TRGSCM.12 | UK | 51.53659 | -0.153415 | *Geotrypetes seraphini* | Dermophiidae | CAPE | 2012 | Yes | |
| CAE2 | UK | 51.53659 | -0.153415 | *Geotrypetes seraphini* | Dermophiidae | CAPE | 2012 | Yes | Yes |
| HR1 | UK | 51.53659 | -0.153415 | *Hyperolius riggenbachi* | Hyperoliidae | CAPE | 2013 | Yes | Yes |
| HR5 | UK | 51.53659 | -0.153415 | *Hyperolius riggenbachi* | Hyperoliidae | CAPE | 2013 | Yes | Yes |
| SAKN-5 | South Africa | -28.71981 | 28.923611 | *NA* | Heleophrynidae | CAPE | 2016 | Yes | Yes |
| SAKN-6 | South Africa | -28.71981 | 28.923611 | *NA* | Heleophrynidae | CAPE | 2016 | Yes | Yes |
| MC58 | South Africa | -23.81647 | 30.0238 | *Hadromophryne natalensis* | Heleophrynidae | CAPE | 2008 | Yes | Yes |
| CCB1 | Spain | 39.87533 | 2.856399 | *Alytes muletensis* | Alytidae | CAPE | 2007 | Yes | Yes |
| CCB15 | Mallorca | 39.87533 | 2.856399 | *Alytes obstetricans* | Alytidae | CAPE | 2015 | Yes | Yes |
| DB8-4 | South Africa | -28.760278 | 28.896389 | *NA* | Pyxicephalidae | CAPE | 2016 | Yes | Yes |
| DB8-2 | South Africa | -28.760278 | 28.896389 | *NA* | Pyxicephalidae | CAPE | 2016 | Yes | Yes |
| TF5a1 | Spain | 39.85755444 | 2.837379981 | *Alytes muletensis* | Alytidae | CAPE | 2007 | Yes | Yes |
| SA546 | South Africa | -29.08682 | 23.79704 | *NA* | Amietia | CAPE | 2017 | Yes | Yes |
| SA152 | South Africa | -29.39124 | 29.73853 | *NA* | Amietia | CAPE | 2017 | Yes | Yes |
| SA186 | South Africa | -29.45001 | 29.51483 | *NA* | Amietia | CAPE | 2017 | Yes | Yes |
| SA190 | South Africa | -29.45001 | 29.51483 | *NA* | Amietia | CAPE | 2017 | Yes | Yes |
| SA211 | South Africa | -29.60708 | 29.33532 | *NA* | Amietia | CAPE | 2017 | Yes | Yes |
| SA243 | South Africa | -29.69575 | 29.4129 | *NA* | Amietia | CAPE | 2017 | Yes | Yes |
| SA266 | South Africa | -29.74601 | 29.20829 | *NA* | Amietia | CAPE | 2017 | Yes | Yes |
| SA280 | South Africa | -29.74601 | 29.20829 | *NA* | Amietia | CAPE | 2017 | Yes | Yes |
| SA286 | South Africa | -29.74601 | 29.20829 | *NA* | Amietia | CAPE | 2017 | Yes | Yes |

Table S2: Burrow-Wheeler Aligner (BWA) mapping statistics. Shotgun sequence reads generated from the two Kihansi samples (AC040803_A and AC290703_A2 ) were aligned to the reference *Bd* mitochondrial genome (JEL423). Mapping statistics were generated using the depth function in samtools. Plots were generated with ggplot2 *(45)*.

A

Kihansi sample – AC040803_A

| Contig | Start position | End position | Number of reads | Bases Covered | Coverage | Mean Depth | Mean base quality | Mean mapping quality |
|---|---|---|---|---|---|---|---|---|
| AATT01000349 | 1 | 36116 | 4435 | 35034 | 97.00 | 8.90 | 39.7 | 56.5 |
| AATT01000350 | 36117 | 57193 | 2404 | 17871 | 84.79 | 8.11 | 39.7 | 57 |
| AATT01000351 | 57194 | 174115 | 14811 | 111560 | 95.41 | 9.14 | 39.7 | 58.8 |

B

Kihansi sample – AC290703_A2

| Contig | Start position | End position | Number of reads | Bases Covered | Coverage | Mean Depth | Mean base quality | Mean mapping quality |
|---|---|---|---|---|---|---|---|---|
| AATT01000349 | 1 | 36116 | 2349 | 33652 | 93.18 | 4.84 | 39.7 | 56.3 |
| AATT01000350 | 36117 | 57193 | 1190 | 17157 | 81.41 | 4.16 | 39.7 | 56.8 |
| AATT01000351 | 57194 | 174115 | 7318 | 107158 | 91.65 | 4.67 | 39.8 | 58.8 |

Table S3: A) Marginal likelihoods and standard deviations (SD) for each BEAST2 relaxed prior demographic models used for phylogenetic dating: Coalescent Bayesian Skyline (CBS), Coalescent Exponential Population (COAL-EXPO), Coalescent Extended Bayesian Skyline (CEBS) and Coalescent Constant Population (COAL-CON). B) Pairwise comparison of the estimate marginal likelihoods given as Bayes Factor according to Kass and Raftery, 1995 *(44)*.

A

| Model | Marginal likelihood | SD |
|---|---|---|
| CBS | -238675.801 | 15.5407585 |
| COAL-EXPO | -238616.7257 | 16.1967686 |
| CEBS | -238472.1813 | 16.0309219 |
| COAL-CON | -238456.2258 | 17.1912247 |

B

| | CBS | COAL-EXPO | CEBS | COAL-CON |
|---|---|---|---|---|
| CBS | 0 | -59.0753 | -203.6197 | -219.5752 |
| COAL-EXPO | 59.0753 | 0 | -144.5444 | -160.4999 |
| CEBS | 203.6197 | 144.5444 | 0 | -15.9555 |
| COAL-CON | 219.5752 | 160.4999 | 15.9555 | 0 |

Table S4: Table representing amphibian species that tested positive with either the pan-lineage *Bd* diagnostic qPCR diagnostic *(31)* or the lineage specific *Bd* qPCR diagnostic *(11)*. Amphibians were swab sampled and metadata recorded across a period of 3 years in the Udzungwa Mountains in Tanzania.

| Species | Bd positive (lineage) | Site | Date | Lat | Long | Altitude (m asl) |
|---|---|---|---|---|---|---|
| Callulina sp. | Positive | Kitolomero | 30/11/2013 | -8.39521 | 35.9824 | 1200 |
| Leptopelis parkeri | Positive (Bd-Cape) | Kaselamgunda | 01/01/2014 | -8.30341 | 35.99532 | 1650 |
| Hyperolius minutissimus | Positive | Kaselamgunda | 05/01/2014 | -8.30341 | 35.99532 | 1650 |
| Hyperolius kihangensis | Positive | Kihanga | 22/02/2014 | -8.3721 | 35.98133 | 1760 |
| Hyperolius kihangensis | Positive (Bd-Cape) | Kihanga | 22/02/2014 | -8.3721 | 35.98133 | 1760 |
| Arthroleptis stenodactylus | Positive | USFR near Kitolomero | 02/03/2014 | NA | NA | 800 |
| Arthroleptis reichei | Positive | Muma | 07/12/2014 | -8.49075 | 35.90381 | 1488 |
| Leptopelis uluguruensis | Positive | Tumbo-Kilola | 17/12/2014 | -8.39103 | 36.0102 | 597 |
| Nectophrynoides tornieri | Positive (Bd-Cape) | Mapepo | 06/01/2015 | -8.52867 | 35.89614 | 742 |
| Amietia angolensis | Positive | Mapepo | 07/01/2015 | -8.52867 | 35.89614 | 742 |
| Amietia angolensis | Positive | Mseo | 30/01/2015 | -8.41295 | 35.92604 | 1597 |
| Phrynobatrachus sp | Positive (Bd-Cape) | Mseo | 08/02/2015 | -8.41295 | 35.92604 | 1597 |
| Hyperolius substriatus | Positive | Mseo | 08/02/2015 | -8.41295 | 35.92604 | 1597 |
| Hyperolius minutissimus | Positive (Bd-Cape) | Mbawala/Site 3 | 10/02/2015 | -8.41293 | 35.94896 | 1647 |
| Arthroleptis xenodactyloides | Positive | Mbawala/Site 3 | 12/02/2015 | -8.41293 | 35.94896 | 1647 |

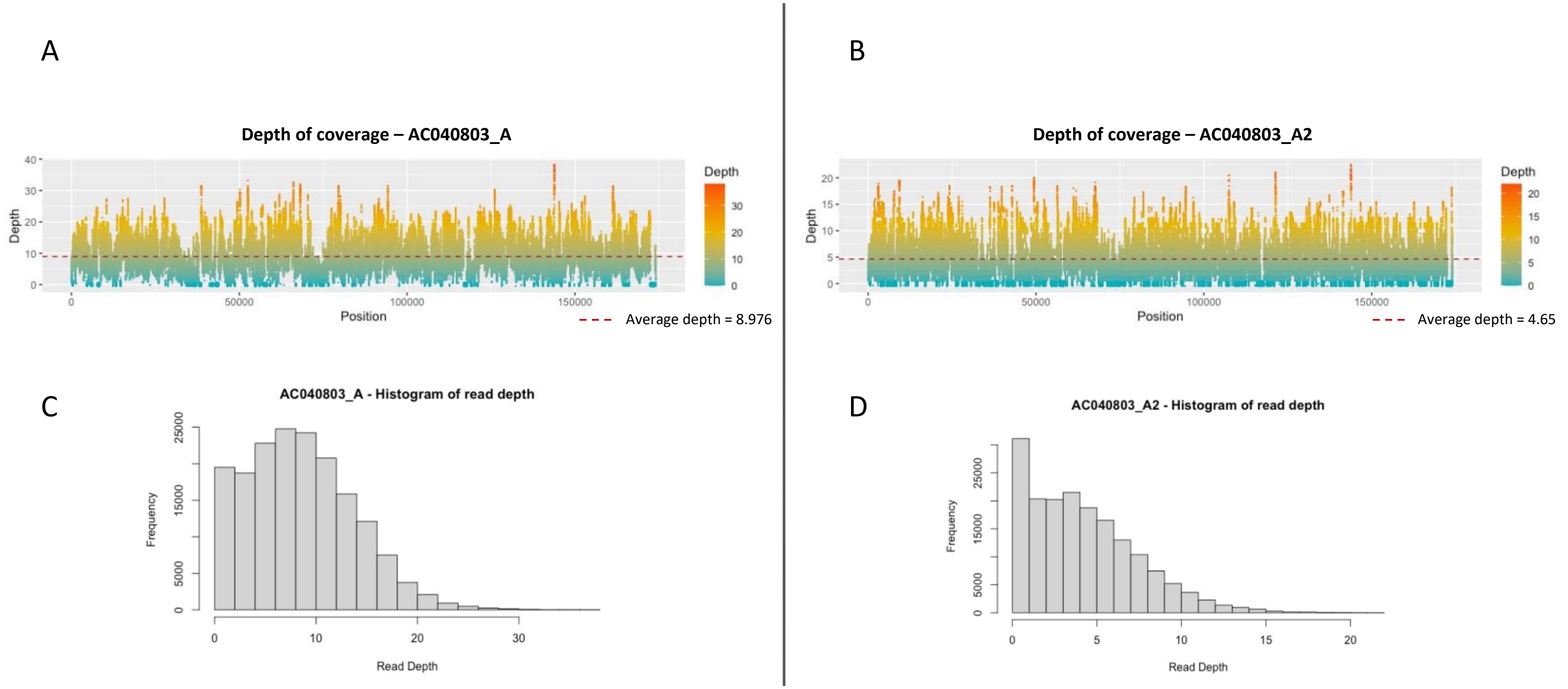


Figure S1: Depth and breadth of coverage - Shotgun sequence reads generated from the two Kihansi samples were aligned to the reference *Bd* mitochondrial genome (JEL423). Depth of coverage was calculated using the depth function in samtools.

A

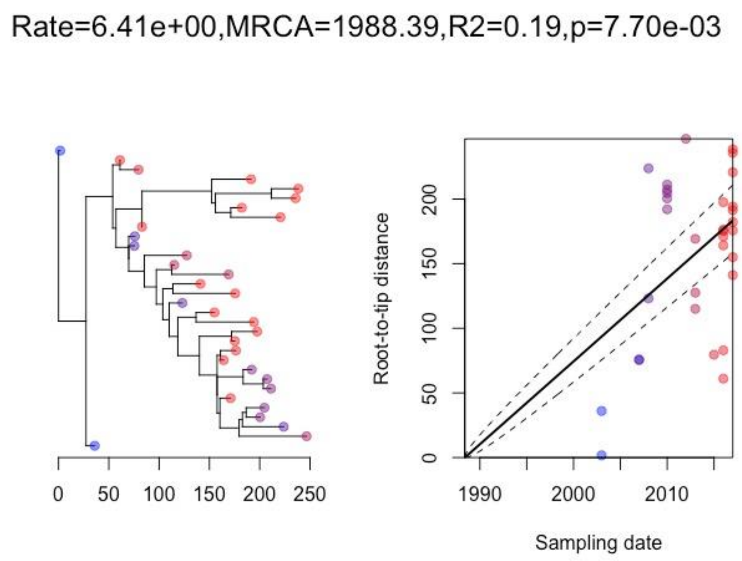


B

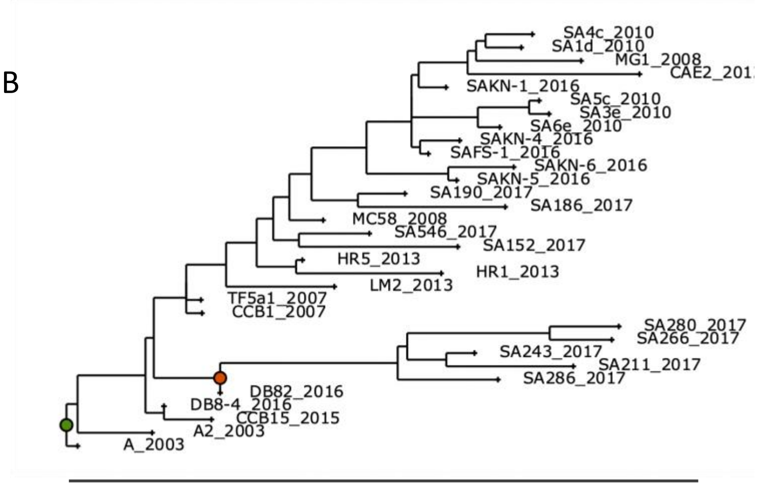


C

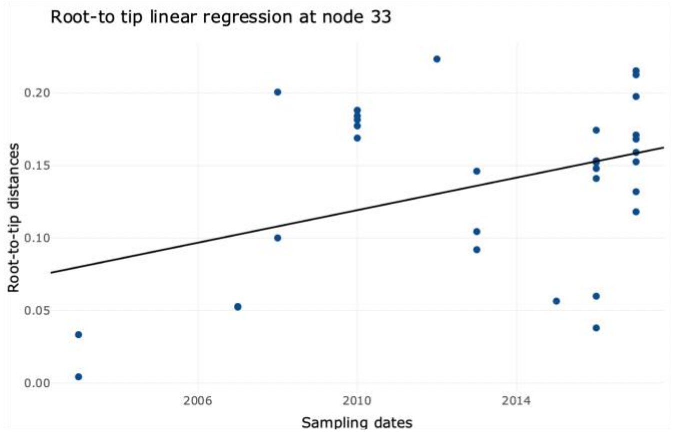


Fig S2. Analysis and confirmation of temporal signal in the pruned mitochondrial alignment of 30 *Bd*CAPE isolates and the two Kihansi samples. A) Output of the BactDating root-to-tip correlation and rate estimation analysis in R. P-value provides significance following 10,000 date randomizations. B) Phylostems analysis which identifies the presence of temporal signal at every evolutionary node of the phylogenetic tree C) Phylostems root-to-tip correlation plot at node 33 (Green node Fig S2B).

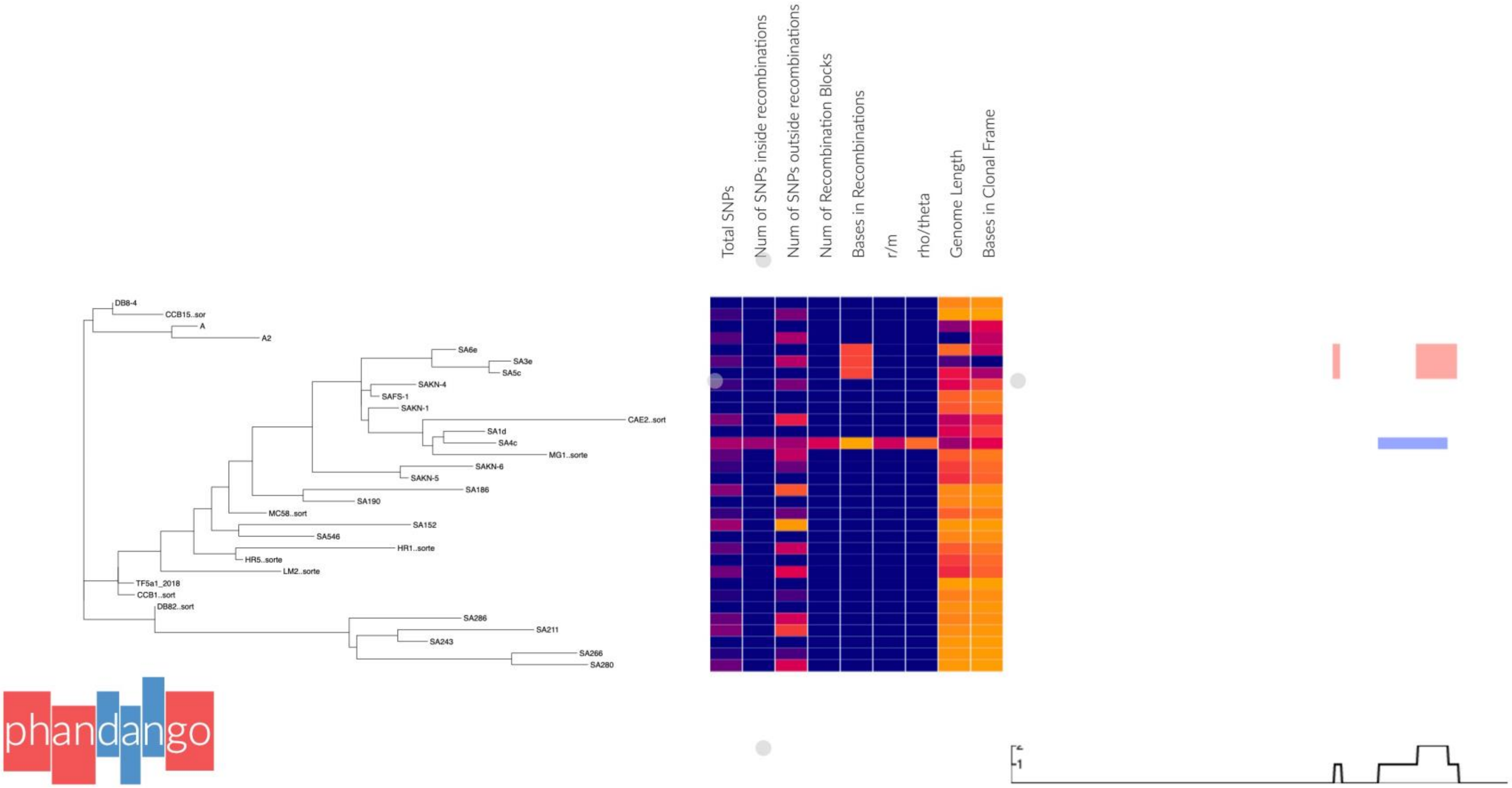


Fig S3. The Identification of putative recombination blocks by distinguishing elevated densities of base substitutions in Gubbins. Visualisation was generated using Phandango. Table insert shows the location of each recombinant block (A-C), the nucleotide coordinates of each block in the mitochondrial alignment (Start – Stop), and the name of the isolates in which that the block is found.

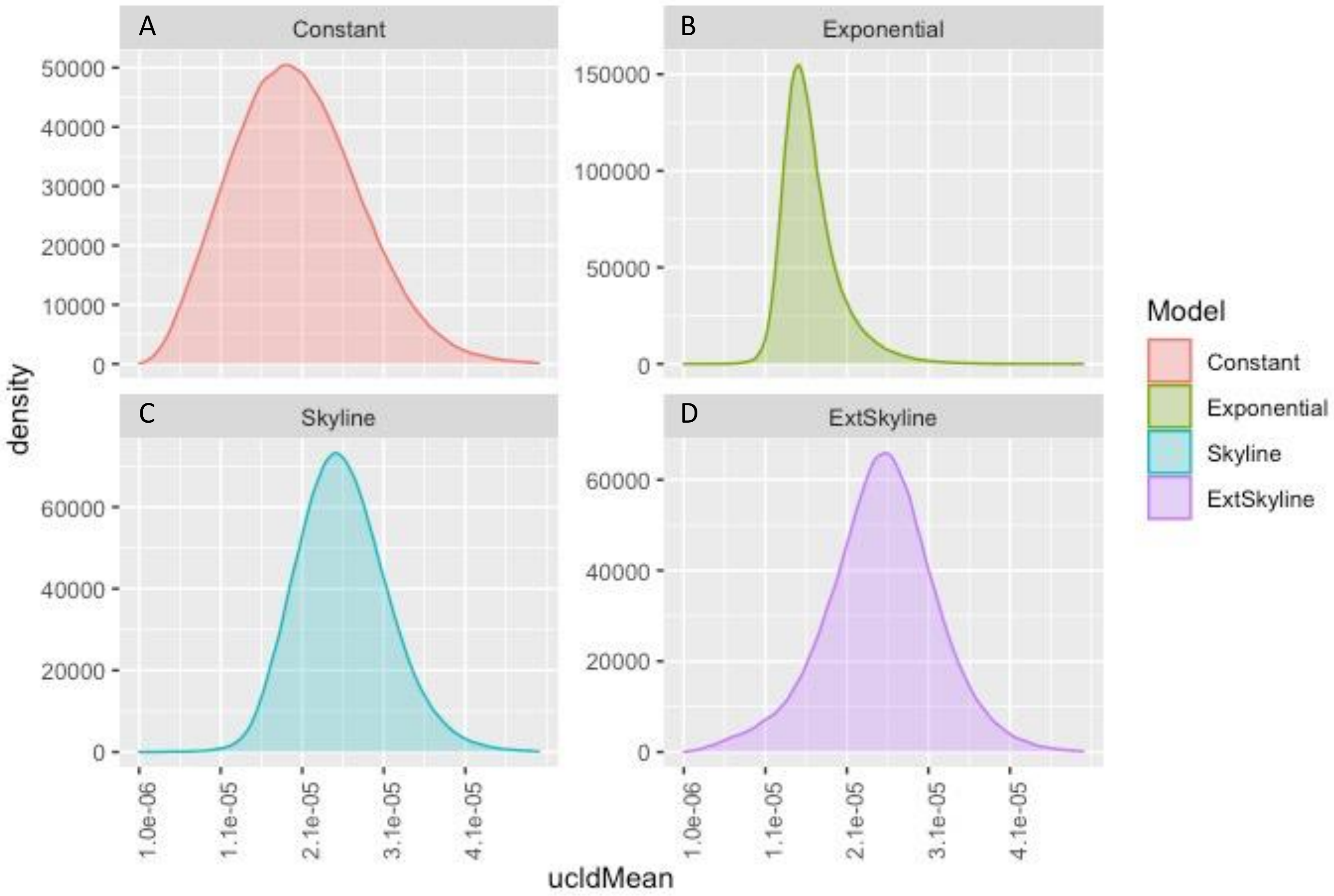


Fig S4. Relaxed clock posterior distributions for inferred clock rates estimated under models specifying four different demographic priors generated using BEAST2: A) Coalescent Constant Population (Constant), B) Coalescent Exponential Population (Exponential), C) Coalescent Bayesian Skyline (Skyline) and D) Coalescent Extended Bayesian Skyline (ExtSkyline).

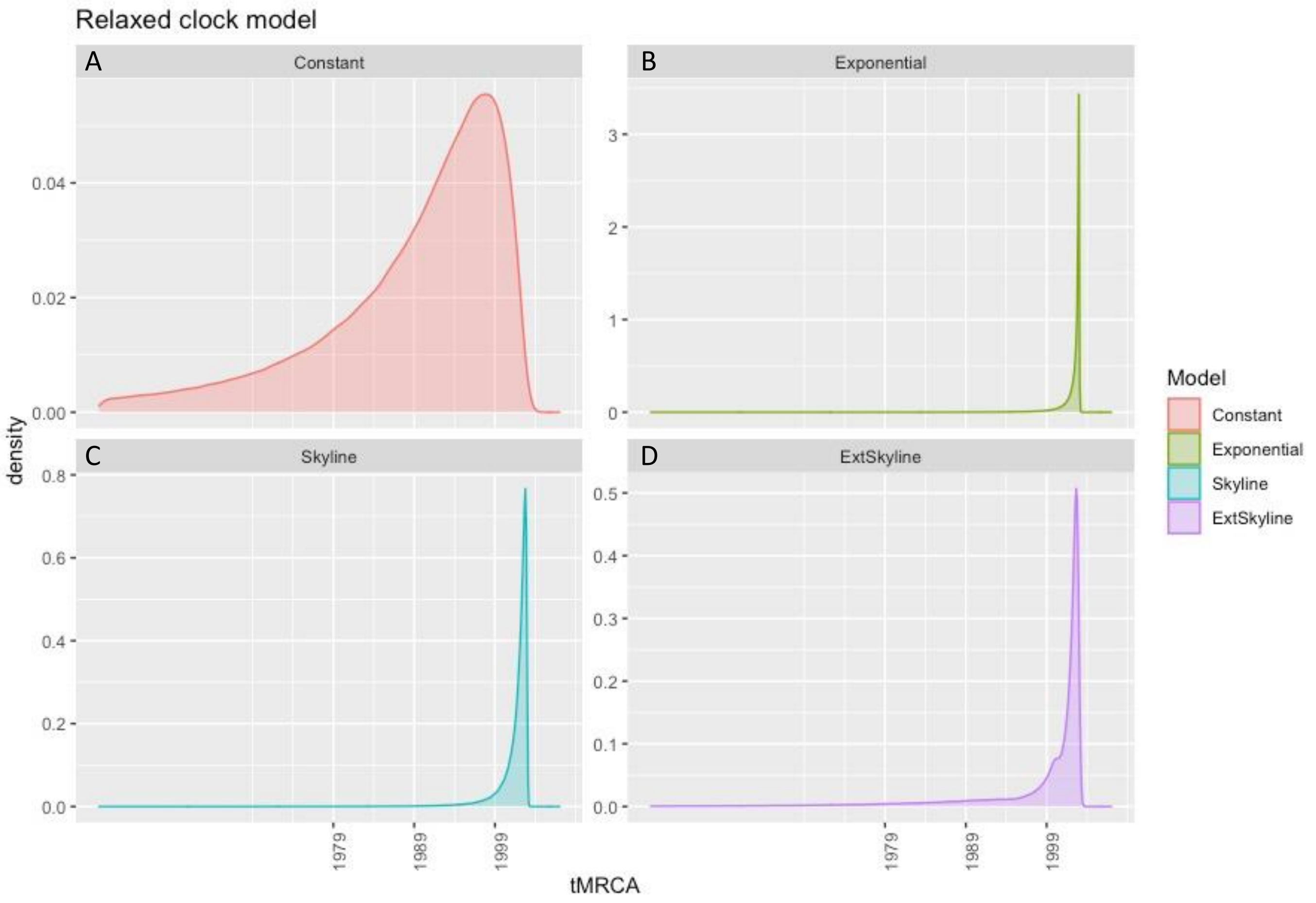


Fig S5. Relaxed clock posterior distributions for inferred tMRCA (tree heights) estimated under models specifying four different demographic priors generated using BEAST2: A) Coalescent Constant Population (Constant), B) Coalescent Exponential Population (Exponential), C) Coalescent Bayesian Skyline (Skyline) and D) Coalescent Extended Bayesian Skyline (ExtSkyline).

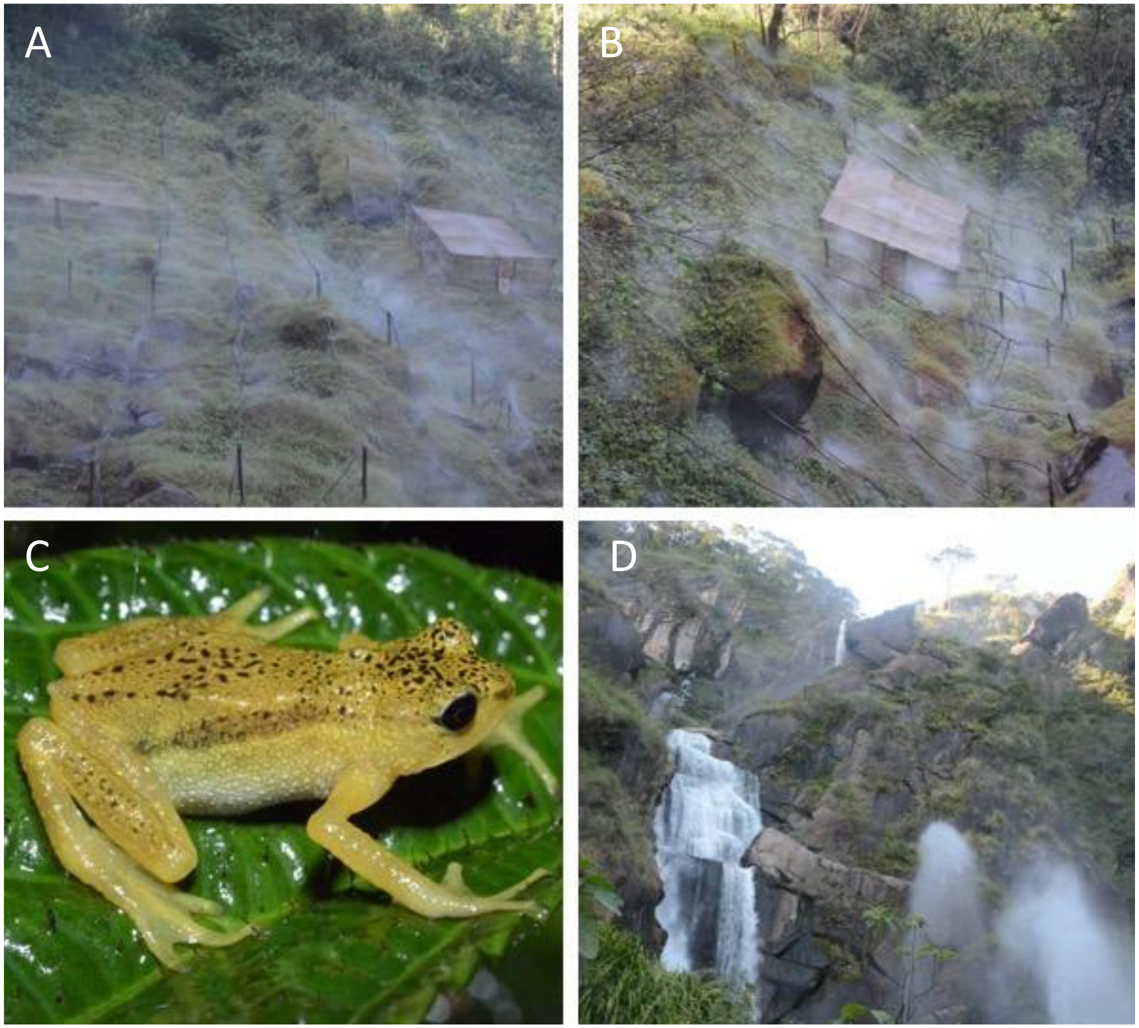


Fig S6. Photographs of the Kihansi spray gorge in the Udzungwa Mountains, Tanzania. A) The upper spray wetlands B) The lower spray wetlands C) A Kihansi spray toad *in situ* D) Waterfalls within the Kihansi gorge.
